# Spatiotemporal control of cell ablation using Ronidazole with Nitroreductase in *Drosophila*

**DOI:** 10.1101/2024.05.05.592574

**Authors:** Gary Teeters, Christina E. Cucolo, Sagar N. Kasar, Melanie I. Worley, Sarah E. Siegrist

## Abstract

The ability to induce cell death in a controlled stereotypic manner has led to the discovery of evolutionary conserved molecules and signaling pathways necessary for tissue growth, repair, and regeneration. Here we report the development of a new method to genetically induce cell death in a controlled stereotypic manner in *Drosophila*. This method has advantages over other current methods and relies on expression of the *E. coli* enzyme Nitroreductase (NTR) with exogenous application of the nitroimidazole prodrug, Ronidazole. NTR expression is controlled spatially using the GAL4/UAS system while temporal control of cell death is achieved through timed feeding of Ronidazole supplied in the diet. In cells expressing NTR, Ronidazole is converted to a toxic substance inducing DNA damage and cell death. Caspase cell death is achieved in a range of NTR-expressing cell types with Ronidazole feeding, including epithelial, neurons, and glia. Removing Ronidazole from the diet restores cell death to normal unperturbed levels. Unlike other genetic ablation methods, temporal control is achieved through feeding not temperature, circumventing developmental complications associated with temperature changes. Ronidazole-NTR also requires only two transgenes, a GAL4 driver and *UAS-NTR*, which is generated as a GFP-NTR fusion allowing for easy setup of large-scale screening of *UAS-RNAi* lines. Altogether, Ronidazole-NTR provides a new streamlined method for inducing cell death in *Drosophila* with temperature-independent ON/OFF control.

## Introduction

Developing methods to eliminate cells has proven key in revealing many of the highly conserved basic biological processes across the animal kingdom. For example, cellular plasticity and transdetermination was demonstrated using fragments of dissected *Drosophila* imaginal tissue transplanted into adult abdomens (Hadorn et al., 1959). Genes controlling growth and cell competition were uncovered through mitotic recombination between wildtype and cell lethal bearing chromosomes (Morata & Ripoll, 1975; Hariharan & Bilder 2006; Marygold et al., 2007; Wang & Dahmann, 2020). Even radiation resulting in double strand breaks and cell death has provided insight into understanding developmental timing and repair mechanisms (Wichmann et al., 2006; Halme et al., 2010; Verghese & Su, 2018). Thus, a wide array of methods exists to induce cell loss and these methods have collectively aided in our understanding of tissue formation, growth, and even regeneration.

To eliminate cells in many model systems including *Drosophila*, one of two approaches is generally used. The first is genetics, where, for example a tissue-specific Gal4 driver controls expression of a UAS-pro-apoptotic transgene gene with the goal of inducing cell death (Grether et al., 1995; P. Chen et al., 1996; White et al., 1996; Wing et al., 1999). Timing of UAS-transgene expression can be controlled by expression of temperature sensitive (ts) Gal80 or auxin-degradable (AD) GAL80 (McGuire et al., 2003; McClure et al., 2022). When GAL80 is active (18°C for GAL80ts or absence of auxin for GAL80AD), the UAS-transgene is not expressed and when Gal80 is inactive (30°C for GAL80ts or presence of auxin for GAL80AD) the UAS-transgene is expressed. On paper, the GAL4/UAS/GAL80 system has the components needed to ablate cells in a spatial and temporal manner, however the system requires three transgenes and GAL80 ON/OFF control takes 12-24 hours in either direction depending on cell and tissue type. Furthermore, changing temperature (18°C to 30°C to 18°C) disrupts animal growth and developmental timing, and it remains unclear whether auxin can be sufficiently flushed from the system to allow UAS-pro-apoptotic transgene expression to be toggled between ON/OFF states (Azevedo et al., 2002; Angilletta et al., 2004; McDonald et al., 2018). The second approach commonly used to eliminate cells and incur tissue damage, include physical methods and depending on the question or tissue of interest range from simple punctures and cuts to force regeneration (Schubiger, 1971; French et al., 1976; Galko & Krasnow, 2004). Physical methods have the advantage of providing great temporal control, however the injuries induced are not cell type specific and the damage induced varies from animal to animal and day to day.

To improve upon current available methods, we developed Nitroreductase for use in *Drosophila*. This method is widely used in Zebrafish and relies on expression of the *E. coli* enzyme, Nitroreductase (NTR) (Drabek et al., 1997; Curado et al., 2007). When NTR expressing cells are exposed to nitroimidazole prodrug, NTR converts the prodrug into a cytotoxic compound leading to DNA damage and cell death. Since its first inception, two major modifications have been implemented. First, is usage of other nitroimidazole derivatives, including Ronidazole and Nifurpirinol, both of which can be used at lower concentrations compared to the first used derivative Metronidazole (Bergemann et al., 2018; Y. Chen et al., 2023). A second modification is a change in the Nitroreductase sequence (T41Q;N71S;F124T) that results in increased catalytic activity (Mathias et al., 2014). The triple NTR mutation (T41Q;N71S;F124T) paired with Ronidazole or Nifurpirinol can effectively kill cells in Zebrafish in just four hours (Bergemann et al., 2018; Y. Chen et al., 2023).

Here, we report the development of *UAS-EmGFP-NTR* transgenic *Drosophila*. Using this transgene in conjunction with tissue specific GAL4 drivers, we find that NTR expressing cells can be effectively eliminated after feeding animals the nitroimidazole prodrug Ronidazole. We use a range of GAL4 drivers and report that this method is effective at killing epithelial cells, neurons, and glia. This method relies on only two transgenes: the tissue specific GAL4 and the *UAS-EmGFP-NTR*. Spatial control is determined by the GAL4 line, whereas temporal control is provided through timed feedings. This method is reversible: prodrug fed animals can be switched to no prodrug food and levels of cell death returned to baseline. As observed with other GAL4/UAS-cell ablation systems, we also report increased Myc and Jun-kinase activity in NTR expressing epithelium, indicative of the injury and regenerative response (Smith-Bolton et al., 2009; Bergantiños et al., 2010; Worley & Hariharan 2022). We have multiple insertions with at least one on each of the major chromosomes. Together, NTR provides a new method to control cell death genetically in *Drosophila* and importantly works in a temperature-independent manner.

## Results

### Ronidazole treatment effectively induces tissue damage in animals expressing Nitroreductase

To determine whether the *E. Coli* enzyme, Nitroreductase (NTR), could cause cell death in the presence of nitroimidazole prodrug in *Drosophila*, we generated *UAS-3xEmGFP-NTR* transgenic animals (Figure 1A and Figure 1-figure supplement 1). The NTR sequence used contains the three previously described point mutations (T41Q;N71S;F124T) and was cloned in frame downstream of three copies of Emerald GFP to facilitate screening (Ilagan et al., 2010; Mathias et al., 2014). To determine whether this method could effectively kill cells, we used *rotundGAL4* (*rnGAL4*) to drive expression of *UAS-3xEmGFP-NTR* in the wing pouch (Kerridge & Thomas-Cavallin, 1988). Cells in the wing pouch give rise to the adult wing structure and when *UAS-reaper* or *UAS-eiger* is expressed with *rnGAL4* in mature third instar larvae, adults eclose from their pupal cases without wings (Smith-Bolton et al., 2009; Harris et al., 2016). We raised *rnGAL4, UAS-3xEmGFP-NTR* transgenic animals on standard Bloomington fly food (Figure 1B). At 72 hours ALH (after larval hatching), animals were transferred to new flood plates containing Bloomington (BL) food mixed with blue dye and different concentrations of each of the three prodrugs, Metronidazole, Nifurpirinol, and Ronidazole. Ingestion of prodrug food was confirmed through visualization of blue dye in the larval digestive tract. After 24 hours of prodrug feeding, animals were transferred back to a regular Bloomington diet and left until adult stages. After adults eclosed, wings were scored and binned into one of two groups, normal or defective (Figure 1B). Animals consuming Ronidazole during development gave rise to adults with defective wings, whereas Metronidazole or Nifurpirinol treated animals gave rise to adults with normal wings only (Figure 1C-E). Furthermore, increasing Ronidazole concentration from 1X (1mM) to 5X (5mM) resulted in increased wing defects in adults (Figure 1E). This suggests that Nitroreductase (NTR) expression in the presence of Ronidazole is effective at killing cells and inducing tissue damage in *Drosophila*.

**Figure 1:**
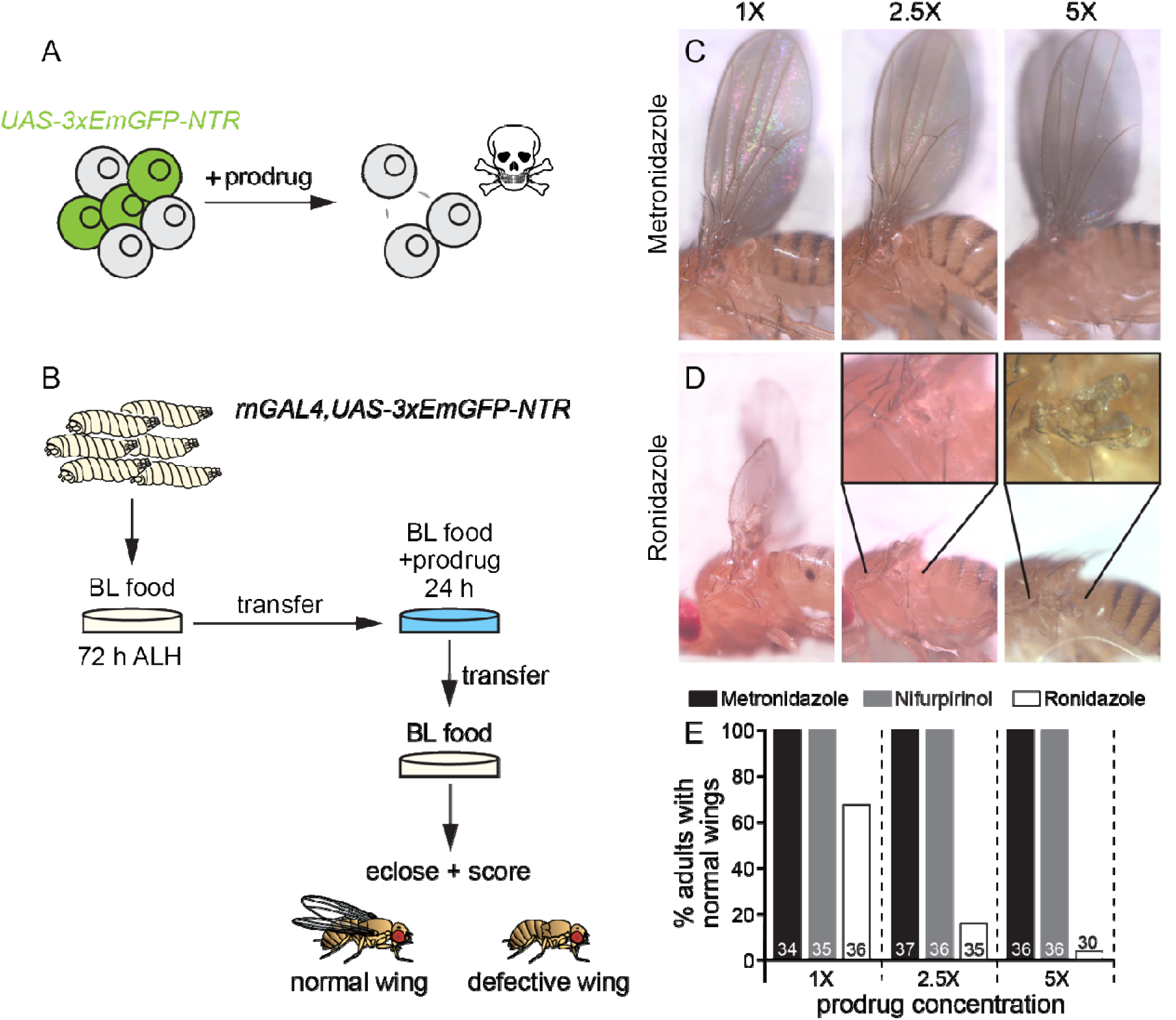
Wing ablation using Nitroreductase (NTR) with Ronidazole. (A) Schematic of tissue specific expression of *UAS-3xEmGFP-NTR* before and after prodrug treatment. (B) Experimental layout with timing of prodrug feeding and screening method to detect cell loss and adult wing phenotypes. (C,D) Adult wing phenotypes of animals expressing NTR in the wing pouch (*rnGAL4*) after treatment with Metronidazole (1X: 4mM, 2.5X:10mM, 5X:20mM) or Ronidazole (1X:1mM, 2.5X:2.5mM, 5X:5mM). (E) Percentage of adult flies with normal wings after prodrug feeding, number of adults scored indicated as number in each bar of the histogram. Only Ronidazole treated animals displayed wing defects. h: hours and ALH: after larval hatching.

To further support this conclusion, we tested two other insertion lines, one located on the X chromosome and one on the second. We again used *rnGAL4* to induce transgene expression and fed larvae 3X (3mM) Ronidazole as depicted in Figure 1B. We used 3X (3mM) Ronidazole in this and all subsequent experiments because 3X (3mM) had less lethality compared to 5X (5mM) and 3X (3mM) induced significant wing defects including complete wing ablations in adult animals (Figure 2 and Figure 2-figure supplement 1). Wing phenotypes after Ronidazole treatment were scored and binned into one of four categories: normal, notched/mis-patterned, small (less than 50% normal size), and no wings (Figure 2A-D). All three lines, X, II, and III (used in Figure 1), when exposed to 3X (3mM) Ronidazole resulted in adults with wings representative of all four classes (Figure 2E). In contrast, when control animals (*rnGAL4* or *UAS-3xEmGFP-NTR* alone) were treated with 3X (3mM) Ronidazole, wing defects were not observed (Figure 2E). Next, we combined insertion lines on the second (II) and third chromosome (III) and found a significant increase in the percentage of adults with wing defects when using 1X (1mM) Ronidazole (compared with Figure 1E). Thus, Ronidazole is effective at ablating NTR-expressing wing imaginal epithelial tissue and the extent of tissue damage is dependent on both the levels of NTR expression and pro-drug Ronidazole treatment.

**Figure 2:**
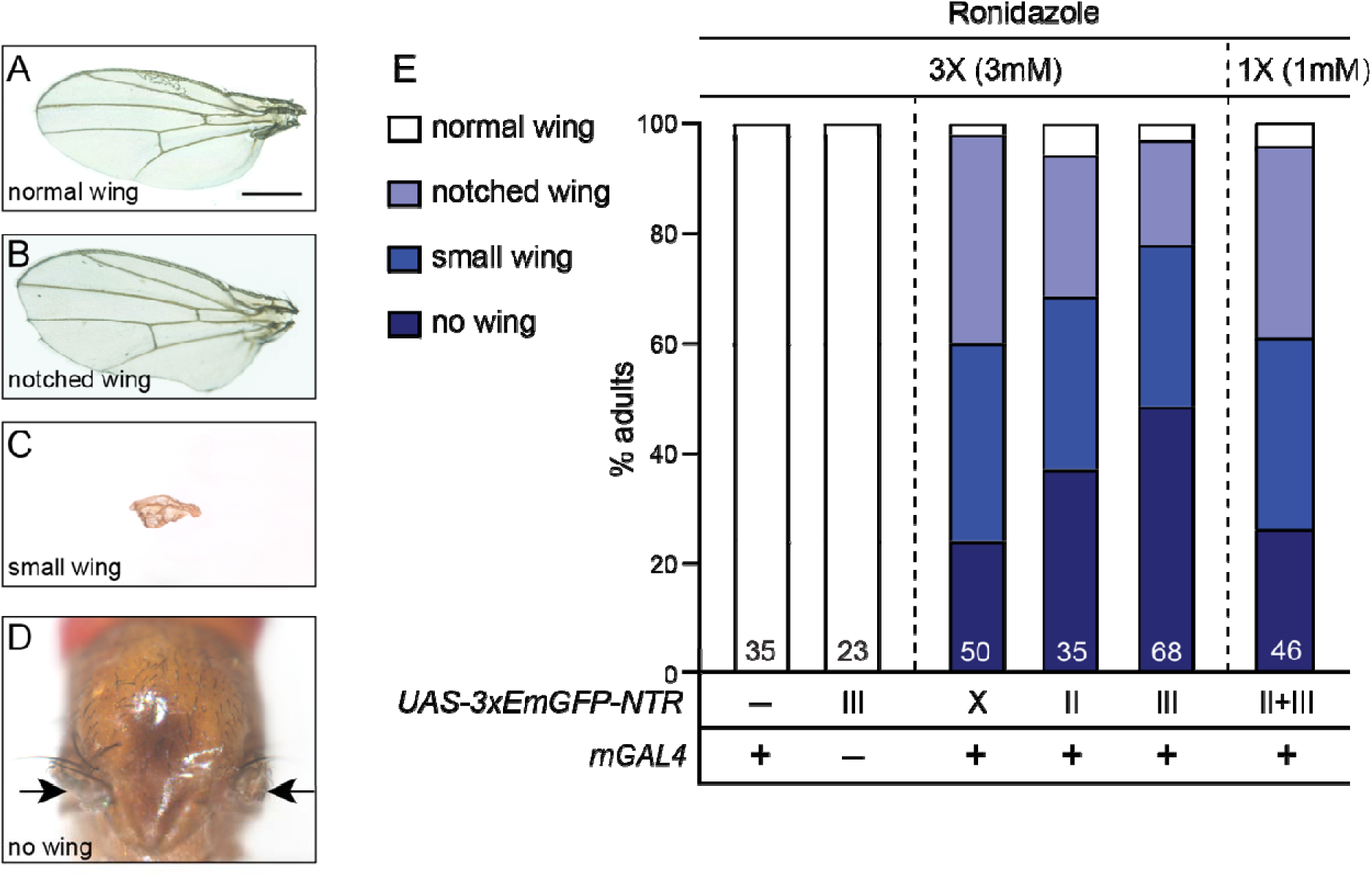
NTR insertion lines on each of the major chromosomes cause wing ablation in Ronidazole treated animals. (A-D) Ronidazole treated NTR animals illicit a range of wing phenotypes from normal (A), to notched or pattern defective (B), to small (C), and no wing (D). (E) Percentages of adults displaying the range of phenotypes categorized based on chromosomal insertion (X, II, or III). Number of adults scored indicated as number in each bar of the histogram. Minus sign indicates absence of indicated transgene (left), plus sign indicates transgene presence. Prodrug concentration listed at top. Scale bar equals 1mM.

### NTR-expressing cells undergo caspase-dependent cell death in the presence of Ronidazole

Next, we assayed cell death of NTR-expressing cells in the presence of Ronidazole. *rnGAL4*, *UAS-3XEmGFP-NTR* larvae were transferred at 72 hours ALH to Ronidazole containing food or no prodrug food as a control (Figure 3A). Larvae were removed from food in four-hour increments and wing imaginal tissues were fixed and stained with antibodies to detect cleaved-Death caspase-1 (Dcp-1) and GFP to label NTR-expressing cells (Figure 3B-K). At 12 hours after prodrug feeding, more than 60% of wing discs showed significant numbers of Dcp-1 positive cells in the GFP expressing wing pouch compared to controls (Figure 3D,I,F,K). At 16 hours, more than 90% of wing discs had substantial numbers of Dcp-1 positive cells (Figure 3E,J,F,K). We conclude that dietary exposure of Ronidazole is effective at inducing caspase-dependent cell death of NTR-expressing cells.

**Figure 3:**
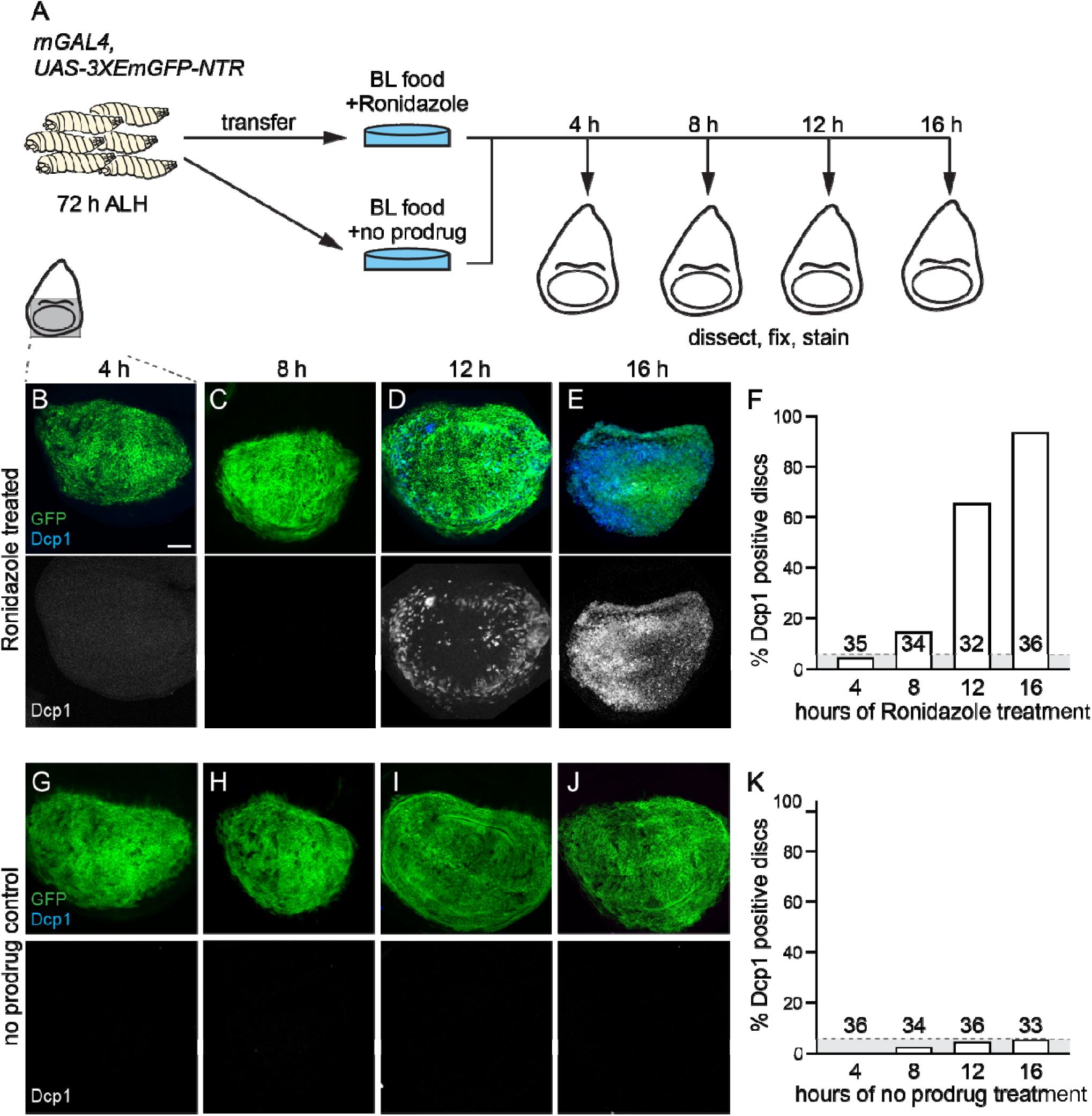
Ronidazole treatment induces caspase-dependent cell death in NTR-expressing animals. (A) Schematic showing timeline of wing imaginal dissection of NTR-expressing animals after hours of prodrug or no prodrug feeding. (B-E) Colored overlay of wing pouch region with grayscale image below of experimental group (*rnGAL4,UAS-3xEmGFP-NTR)* after 4 to 16 hours of Ronidazole feeding. Molecular markers indicated in panels. (F) Percentage of imaginal discs with significant numbers of Dcp-1 positive cells above baseline (grey highlight implemented from control). (G-J) Colored overlay of wing pouch region with grayscale image below of control group (*rnGAL4,UAS-3xEmGFP-NTR)* after 4 to 16 hours of no prodrug feeding. (K) Percentage of imaginal discs showing baseline Dcp-1 staining grey (highlight). Number of adults scored indicated as number in each bar of the histogram. Scale bar equals 40μm. h: hours and ALH: after larval hatching.

### Removing Ronidazole from diet returns cell death levels to baseline in NTR-expressing animals

Next, we asked whether Ronidazole could be effectively removed from the system and cell death levels returned to baseline. First, we determined the time until clearing of blue Ronidazole-containing food from the larval gut. *rnGAL4*,*UAS-3xEmGFP-NTR* larvae were transferred at 72 hours ALH to Ronidazole containing food mixed with blue dye for 12 hours (Figure 4A). After 12 hours, all animals had consumed Ronidazole based on presence of blue within their digestive tract (Figure 4B). Animals were then transferred to BL food with no prodrug and no dye (Figure 4A). After four hours of feeding, approximately 50% of larvae cleared Ronidazole containing food from their gut based on the absence of blue dye and by 12 hours, more than 90% cleared Ronidazole food (Figure 4B).

**Figure 4:**
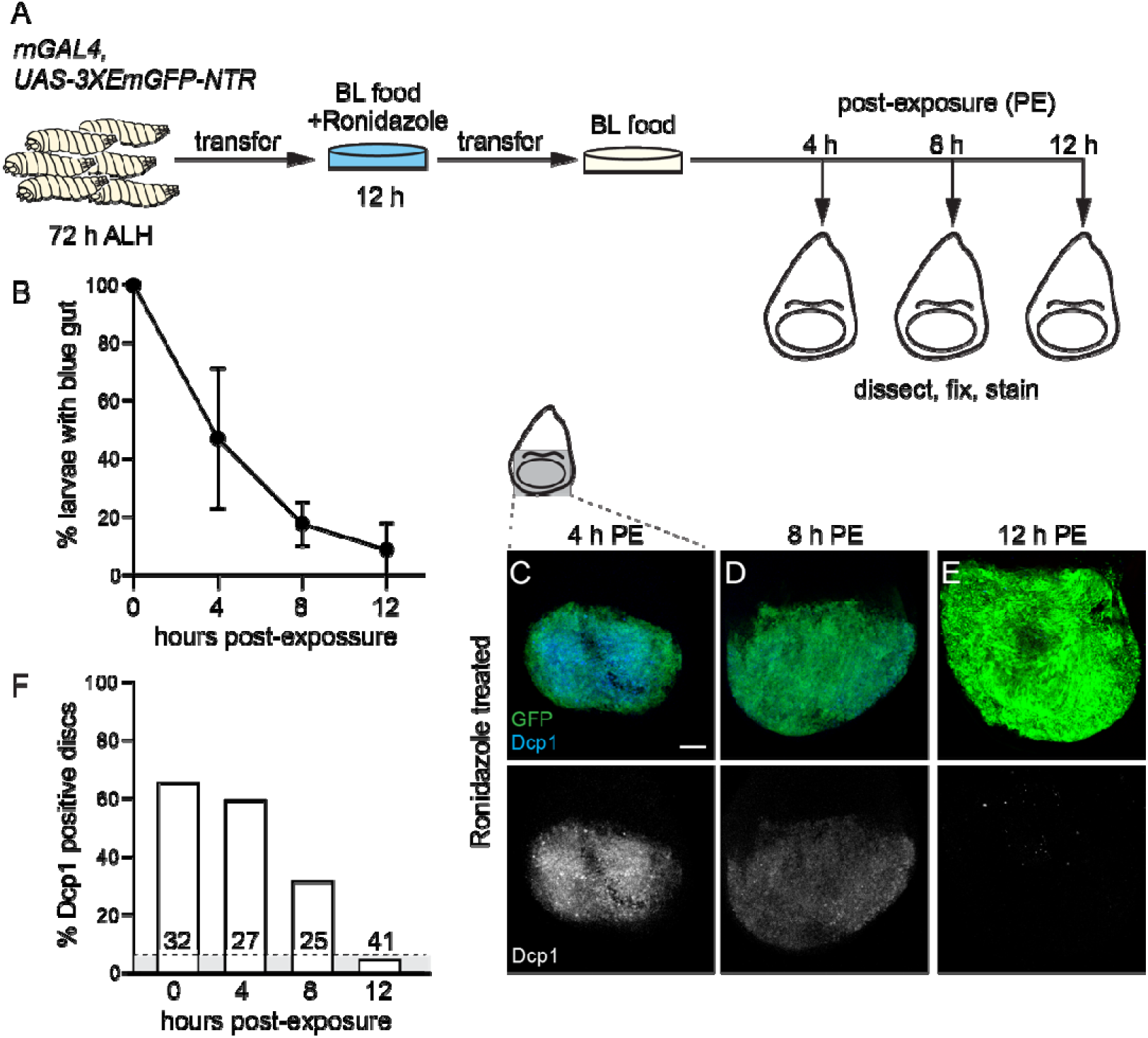
Cell death levels return to baseline after removal of Ronidazole from diet. (A) Schematic showing timeline of wing imaginal dissection after hours of prodrug feeding and then return to no prodrug food of NTR-expressing animals. (B) Percentage of larvae with blue dye remaining in their digestive tract after transfer to no prodrug food without dye. Each data point represents the mean of three experimental groups with each experimental group containing twelve animals. Error bars indicate standard deviation (C-E) Colored overlay of wing pouch region with grayscale image below of experimental group (*rnGAL4,UAS-3xEmGFP-NTR)* after 4 to 12 hours of no prodrug feeding, post-exposure (PE). Molecular markers indicated in panels. (F) Percentage of imaginal discs showing Dcp-1 staining with gray box designating baseline from Figure 3K. Number of animals scored indicated as number in each bar of the histogram. Scale bar equals 40μm. h: hours and ALH: after larval hatching.

Next, we assayed cell death in only the larvae which had cleared blue food from their gut. At four hours after transfer to BL food only (post-Ronidazole exposure), approximately 60% of wing discs still showed significant numbers of Dcp-1 positive cells in the GFP expressing wing pouch (Figure 4C,F). At eight hours post-exposure, approximately 30% of wing discs showed significant numbers of Dcp-1 positive cells and by 12 hours, the percentage was even further reduced (Figure 4D-F). This indicates that Ronidazole remains at biologically active concentrations within the system for less than 12 hours after animals are transferred to food without the prodrug. Furthermore, continuous exposure to prodrug is required to sustain death of NTR-expressing cells.

### Ronidazole treated NTR imaginal tissue elicit an injury and regenerative response

During development, damaged imaginal tissue secrete a Dilp-8 stop signal, which results in a developmental delay allowing time for damaged tissue to self-repair (Colombani et al., 2012; Garelli et al., 2012). We asked whether tissue damaged using the Ronidazole-NTR method would elicit a developmental delay. At 72 hours ALH, *rnGAL4*, *UAS-3xEmGFP-NTR* larvae were transferred to Ronidazole containing food or no prodrug food (Figure 5A). After 24 hours of feeding, animals were then transferred back to BL food without prodrug and time to pupation assayed. We found that NTR expressing larvae treated with Ronidazole took significantly longer to reach pupation compared to the no prodrug fed control or Ronidazole fed *rnGAL4* animals alone (Figure 5B). This suggests that Ronidazole-NTR induced damage is sufficient to induce developmental delay.

**Figure 5:**
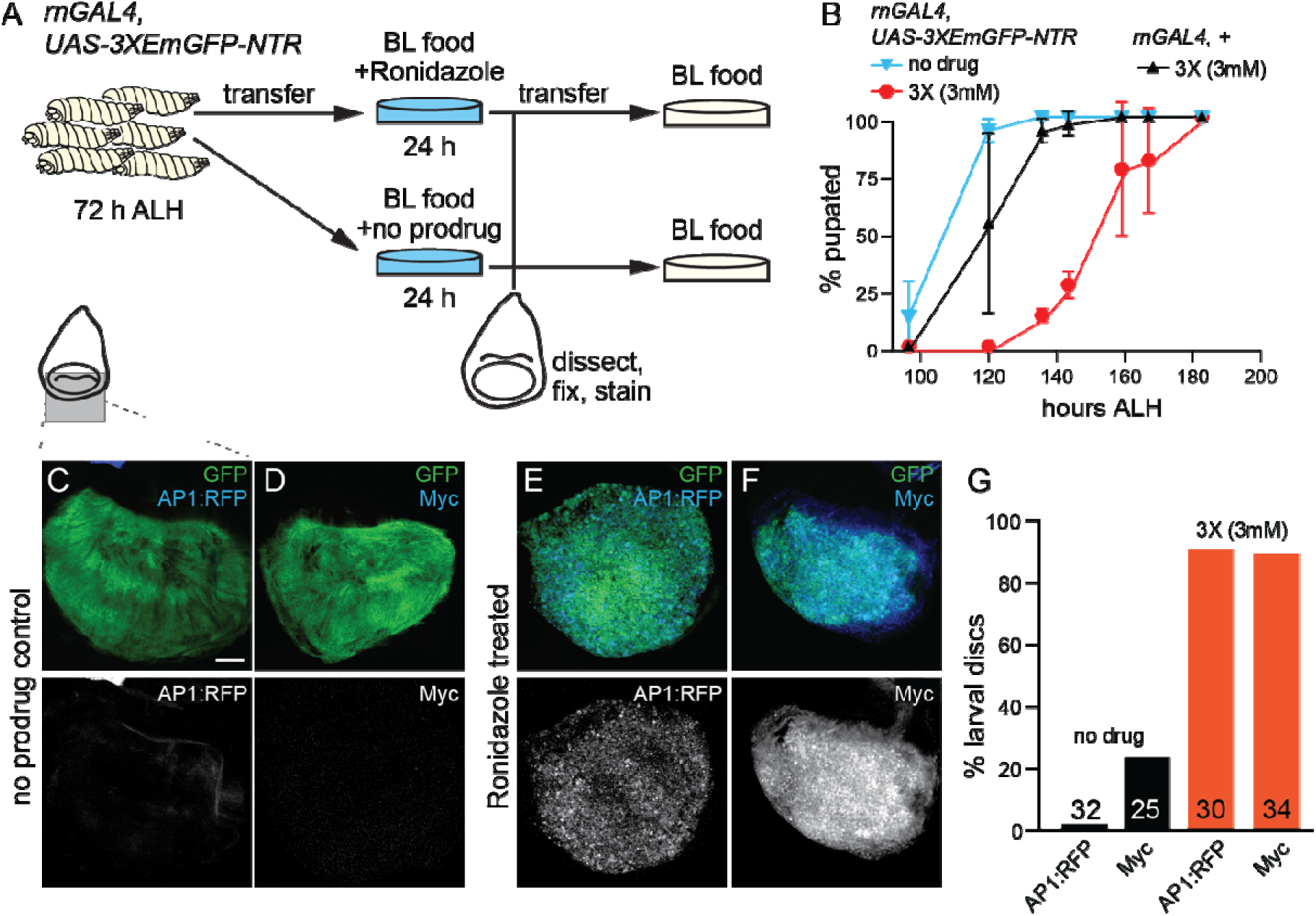
Ronidazole treated NTR-expressing wing discs display hallmarks of the injury and regenerative response. (A) Schematic with timing of prodrug feeding and dissections as well as prodrug feeding and return to no prodrug for assaying developmental timing. (B) Time to pupation from freshly hatched larval stage to P0 white pupal formation between NTR-expressing and non-expressing animals treated with or without prodrug. Each data point represents the mean of three experimental groups with each experimental group containing twelve animals. Error bars equal standard deviation. (C-F) Colored overlay of wing pouch region with grayscale image below. Molecular markers and/or transgenes used indicated in panels. (G) Percentage of imaginal discs showing Myc staining or AP1 reporter expression. Number of discs scored indicated as number in each bar of the histogram. Scale bar equals 40μm. h: hours and ALH: after larval hatching.

Next, we investigated whether molecular markers consistent with an injury and regenerative response are expressed in Ronidazole-NTR treated discs. We used the transcriptional reporter *AP1:RFP* as a readout for JNK pathway activity and antibodies to detect Myc (Chatterjee & Bohmann, 2012). Both are expressed in damaged wing imaginal tissue and both positively regulate the regenerative response (Bosch et al., 2008; Smith-Bolton et al., 2009; Bergantiños et al., 2010; Furrer et al., 2010; Worley et al., 2018). At 72 hours ALH, *rnGAL4*, *UAS-3xEmGFP-NTR* larvae were transferred to Ronidazole containing food or no prodrug food (Figure 5A). After 24 hours of feeding, wing imaginal tissue was dissected and processed for imaging. We found significant increases in the percentage of wing tissue expressing the *AP1:RFP* transcriptional reporter and Myc in Ronidazole treated animals compared to no prodrug controls. Thus, Ronidazole-NTR shares common molecular features with other cell ablation methods and importantly has the added benefit of being controllable in a temperature-independent manner.

### Ronidazole-NTR effectively induces death of both neurons and glia

Finally, we asked whether Ronidazole-NTR could be used to eliminate cells in other tissue types. We used *eyGAL4* to express *UAS-3xEmGFP-NTR* in retinal precursors of the developing eye disc. At seventy two hours ALH, *eyGAL4,UAS-3xEmGFP-NTR* larvae were transferred to Ronidazole containing food or no prodrug food. After 24 hours of feeding, animals were then returned to no prodrug food and eye morphology scored in adults as depicted in Figure 1. We found significant defects in eye shape and head structure in the majority of Ronidazole fed animals compared to the no prodrug fed controls (Figure 6A-C). Next, we assayed cell death and found a significant percentage of eye discs with activated-caspase after 24 hours of Ronidazole feeding whereas control discs had essentially none (Figure 6D-F). Furthermore, activated caspase was restricted to the GFP positive eye field and little to no caspase activity was observed in neighboring presumptive antenna or head capsule tissue.

**Figure 6:**
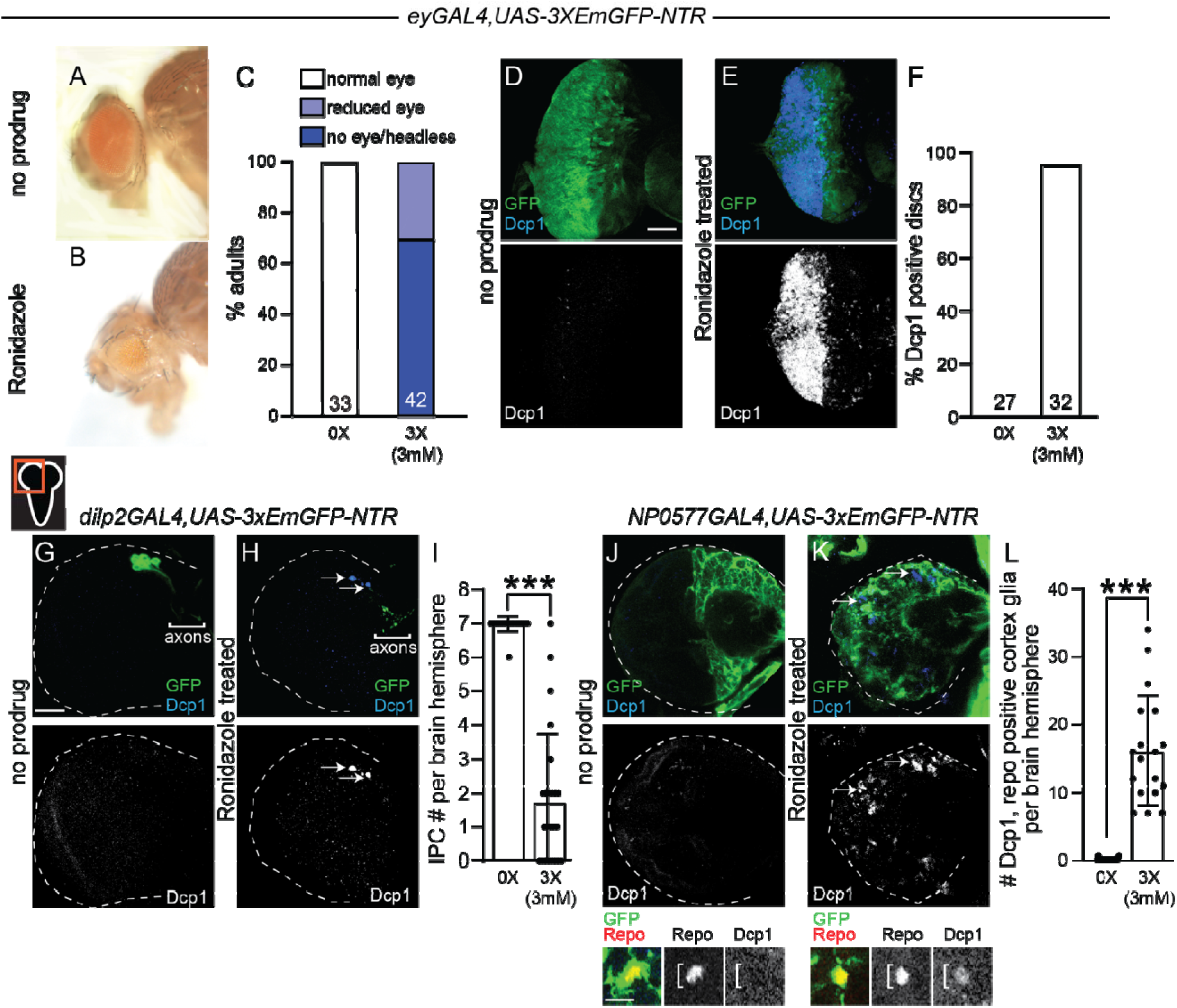
Ronidazole-NTR is effective at killing both neurons and glia. (A,B) Bright field image of *ey-Gal4,UAS-3xEmGFP-NTR* adult heads treated with or without prodrug. (C) Percentage of *ey-Gal4,UAS-3xEmGFP-NTR* adults with eye phenotypes binned into one of three categories. (D,E) Colored overlay of eye imaginal disc with grayscale image below. Molecular markers indicated in panels and genotype above. (F) Percentage of eye imaginal discs with significant numbers of Dcp-1 positive cells. (G,H) Colored overlay of brain hemisphere with grayscale image below. Molecular markers indicated in panels and genotype above. (I) Number of IPCs per brain hemisphere. Each dot represents one brain hemisphere. (J,K) Colored overlay of brain hemisphere with grayscale image below. Molecular markers indicated in panels and genotype above. Below, high magnification image of a cortex glia co-labeled with Repo. (L) Number of Repo positive cortex glia that are Dcp-1 positive. Each dot represents one brain hemisphere. Error bars equal standard deviation. ***p-value<0.0001 (unpaired students T-test). Scale bar equals 40μm except in high magnification panels of J and K, where scale bar equals 10μm. h: hours and ALH: after larval hatching.

Subsequently, we asked whether Ronidazole could cross the blood brain barrier (BBB) and induce death of NTR-expressing neurons and glia. We used *dilp2GAL4* to express *UAS-3xEmGFP-NTR* in the insulin producing cells (IPCs). After 24 hours of Ronidazole treatment, we found a significant number of Dcp-1 positive, GFP positive IPCs in the anterior medial region of each brain hemisphere, which corresponded with a reduction in number of IPCs compared to no prodrug fed controls (Figure 6G-I). Morphology of IPC axons was also disrupted with blebbing observed in Ronidazole treated larvae but not in prodrug controls (Figure 6G-H). Consistent with IPC death, brains of Ronidazole treated animals were smaller compared to no prodrug controls.

Next, we used *NP0577GAL4* to express *UAS-3xEmGFP-NTR* in cortex glia. Cortex glia are a glia subset that ensheath neuroblasts and their newborn progeny. After 24 hours of feeding, we found significant defects in cortex glia morphology based on GFP expression in Ronidazole treated animals compared to no prodrug fed animals (Figure 6J,K). Furthermore, we found a significant number of Dcp-1 positive, GFP positive, Repo positive cortex glia in Ronidazole treated animals (Figure 6J,K bottom panels and quantified in L). We conclude that Ronidazole crosses the BBB and is effective at killing both NTR-expressing neurons as well as NTR-expressing glia.

## Discussion

Here we report the development of a new method to genetically eliminate cells in *Drosophila* that is temperature-independent. This method requires the administration of a nitroimidazole prodrug in the presence of the enzyme Nitroreductase (NTR). The prodrug-NTR method is widely used in other model systems, yet until now had not been developed and tested in *Drosophila* (Clark et al., 1997; Curado et al., 2007; Y.-C. Chen et al., 2023). One clear advantage in using this method over others in *Drosophila* is its means for temporal control, drug feeding versus temperature shifts. It is widely appreciated that temperature shifts from 18°C to 30°C in *Drosophila* globally affect a range of basic cellular processes including transcription and control of growth and proliferation rates, many times in ways that are difficult to predict (McCabe & Partridge, 1997; Azevedo et al., 2002; McDonald et al., 2018). These effects due to temperature change can obfuscate and sometimes even mask phenotypes of cells and tissues responding to damage during normal development and regeneration (Ryoo and Bergmann, 2012; Herrera et al., 2013; Myat et al., 2019).

Recently, a number of new genetic methods have been developed to control cell elimination in a mostly temperature-independent manner. These include GAL80-auxin as well as the DUAL control method (McClure et al., 2022; Zirin et al., 2024; Harris *et al.,* 2020). Using GAL80-auxin, induction of UAS-transgene expression occurs after 18 hours of auxin feeding (McClure *et al.,* 2022), whereas in the DUAL control system LexAop-transgene expression is induced within six hours, however achieving the desired degree of tissue ablation can be limited by the toxicity caused by long heat shocks (Harris *et al.,* 2020). Moreover, neither of these methods have been shown to work in cell types within the brain. With the Ronidazole-NTR method, induction of cell death occurs within twelve hours after prodrug feeding and cell death levels return to baseline twelve hours later simply by removing prodrug from diet. No temperature shift or heat-shock are needed to activate the prodrug-NTR system, and the duration of the ablation period can easily be adjusted by changing the duration of pro-drug exposure. Importantly, the Ronidazole-NTR method is effective at inducing cell death across multiple different cell types and only requires two transgenes (whereas GAL80-auxin and DUAL control require at least three). In addition, because the *UAS-EmGFP-NTR* transgene is a GFP fusion, large scale screening of *UAS-RNAi* lines with subsequent screening of phenotypes based on GFP is easily achievable. Nevertheless, we anticipate that prodrug-NTR method in *Drosophila* could still be improved. A codon optimized version of NTR for *Drosophila* could enhance enzyme kinetics allowing for shorter prodrug treatments at reduced concentrations (Zhao et al., 2017). In addition, Nitroreductase isolated from *Vibrio vulnificus* was recently shown to enhance cell death with Metronidazole in Zebrafish (Sharrock et al., 2022). Developing methods to apply Ronidazole to cells and tissues during nonfeeding stages could also be useful. In conclusion, the Ronidazole-NTR method provides an additional means to conditionally eliminate defined cell populations in *Drosophila* and offers clear advantages over other current available methods.

## Materials and Methods

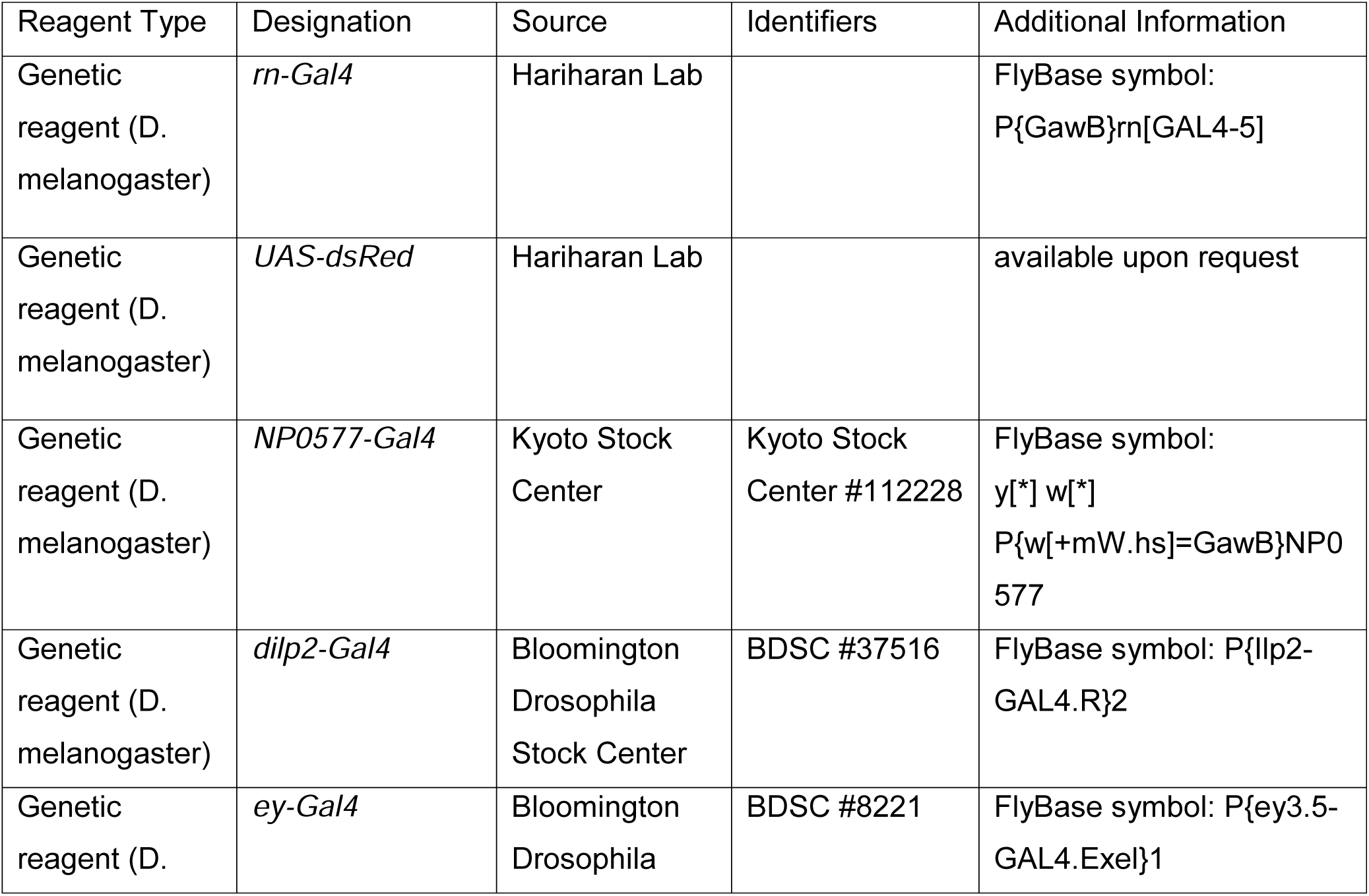
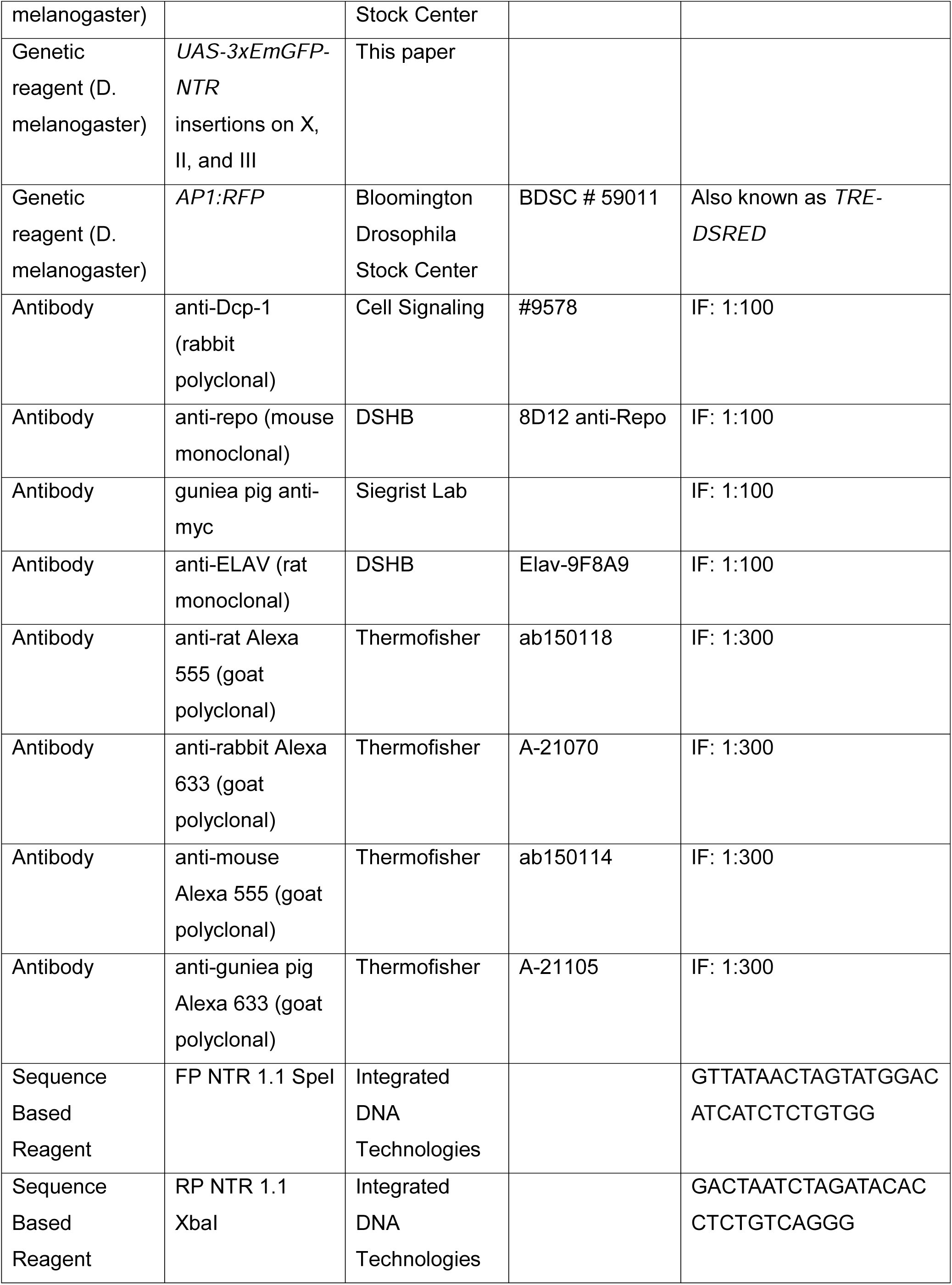
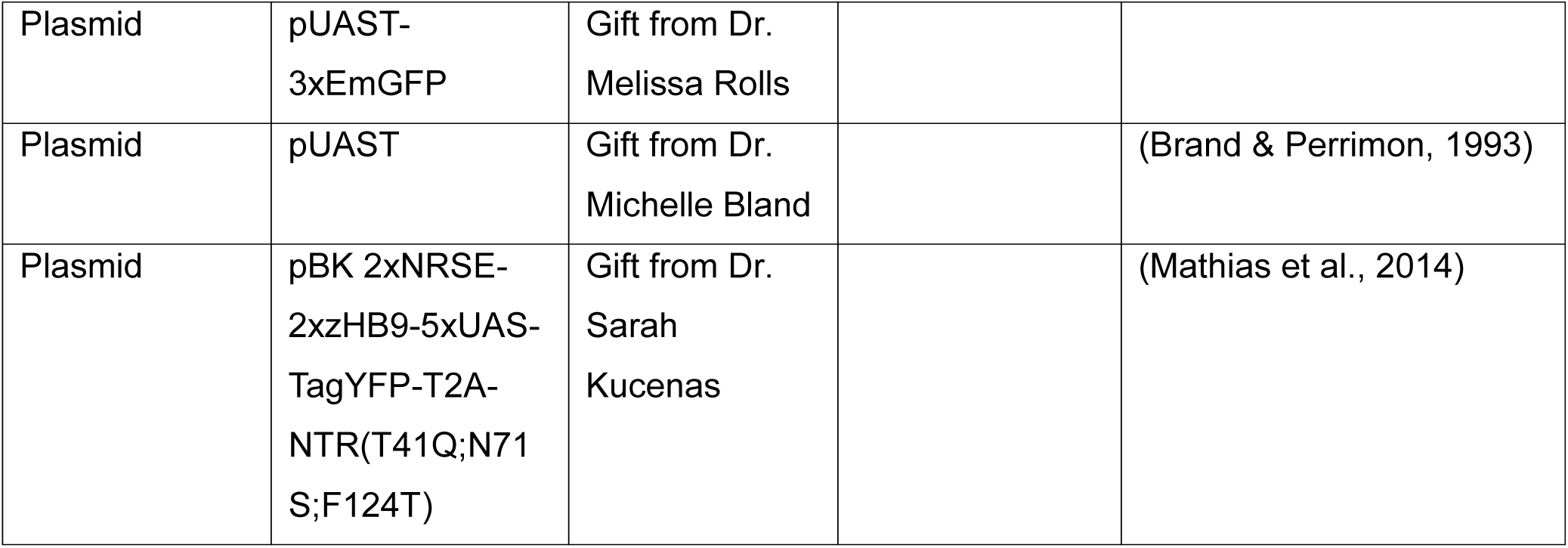

### Fly Rearing, Stocks, and Genetics

All animals were raised in uncrowded conditions at 25°C on a 24-hour light dark cycle (12 on/12 off) on a Bloomington fly food diet (BL) commercially produced by Archon Scientific (Corn Syrup-Soy media W1).

### Plasmid construction and transgenesis

The Nitroreductase (nfsB) sequence was PCR amplified from the plasmid pBK 2xNRSE-2xzHB9-5xUAS-TagYFP-T2A-NTR(T41Q;N71S;F124T) using primer sequence FP: 5’ GTTATAACTAGTATGGACATCATCTCTGTGG 3’ and RP: 5’ GACTAAGGTACCTTACACCTCTGTCAGGG 3’. Restriction enzyme sequences for SpeI and KpnI were included in the FP and RP sequences respectively and cloned into a pUAST plasmid (Brand & Perrimon, 1993) Three copies of Emerald GFP (from pUAST-3xEmGFP, gift from Dr. Melissa Rolls) were placed upstream using EcoRI and SpeI. The final version of the pUAST-3xEmGFP-NTR plasmid was verified through Sanger sequencing before injection. Single embryo injections were carried out by BestGene. Ten independent transgenic lines were recovered each with one insertion on one of the major chromosomes. Three of the ten lines were further characterized and are referenced as X, II, and III.

### Application of prodrug and animal staging

Freshly hatched larvae (zero to one hour after embryonic hatching) were collected and placed onto BL food plates for designated times. Animals were then washed and transferred to BL food containing prodrug with blue dye for designated times. Prodrug concentrations used: Metronidazole 4mM (1X), 10mM (2.5X), 20mM (5X), Nifurpirinol 10uM (1X), 25uM (2.5X), 50uM (5X), and Ronidazole 1mM (1X), 2.5mM (2.5X), 3mM (3X), 5mM (5X). After prodrug feeding, larvae were washed and visually inspected for presence or absence of blue food in gut.

### Tissue Dissections, Antibody Labeling, and Imaging

Wing and eye imaginal tissues were dissected, fixed, and stained following standard methods (Spratford & Kumar, 2014). Brains were dissected and processed as previously described (Sipe & Siegrist, 2017; Pahl et. al, 2019). Primary antibodies used are listed in the Key Resources Table. To detect primary antibodies, Alexa-Fluor conjugated secondary antibodies listed in the Key Resources Table were used. Images were acquired using a Leica SP8 laser scanning confocal microscope equipped with a 63x/1.4 NA or 40x/1.3 NA oil-immersion objectives. Images of adult eyes and wings collected using a Zeiss AxioZoom V16.

### Quantification and image analysis

All images were processed using Fiji and Adobe Photoshop software and figures were assembled using Adobe Illustrator software. Larva and adult drawings were acquired through Scidraw.io. Histograms with percentages represent the mean and histograms displaying cell counts represent mean with standard deviation. Imaginal discs were binned into either positive or negative for Dcp-1, Myc, or *AP1:RFP*, based on number of GFP expressing cells co-expressing the marker of interest. Imaginal discs were binned into the positive group if more than 30 cells were co-labeled with GFP and marker of interest, negative if less than. Statistical significance was determined using unpaired two-tailed Student’s t-tests in Prism 10.

## Acknowledgments

We thank Iswar Hariharan, Melissa Rolls, the Bloomington *Drosophila* Stock Center, the Kyoto stock center and the Developmental Studies Hybridoma Bank for providing plasmids, flies, and antibody reagents. We thank all Siegrist and Worley lab members for providing helpful comments and suggestions on the manuscript and research project.

## Figure Legends

**Figure 1-Figure Supplement 1.**
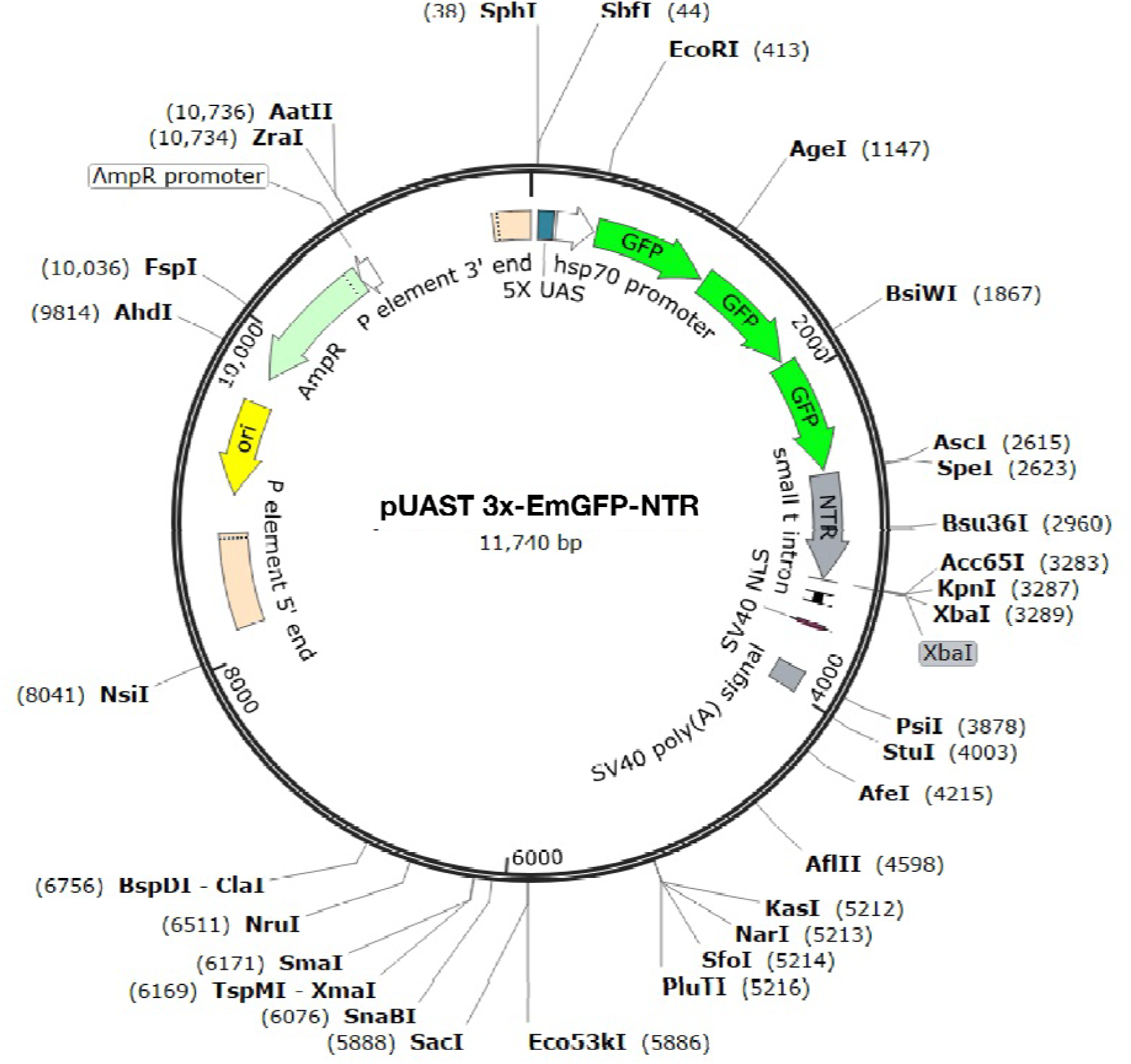
Sequence of pUAST-3xEmGFP-NTR. A genetic map of the pUAST plasmid with 3XEmGFP-NTR inserted within.

**Figure 2-Figure Supplement 1.**
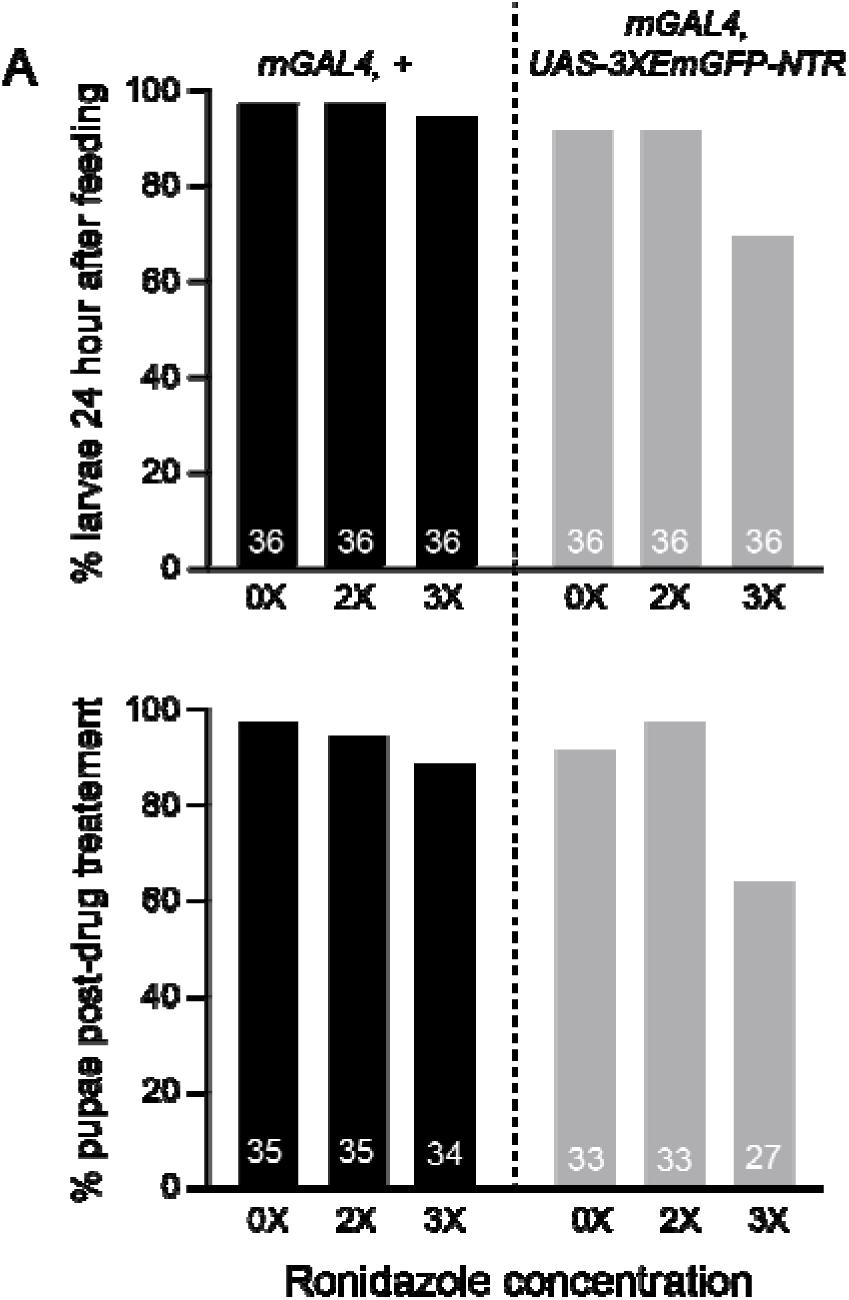
Effects of Ronidazole treatment on survival of control and NTR-expressing animals. (A) Top panel, percentage of larva that survive after 24 hours of no prodrug food versus Ronidazole food and bottom panel, percentage of these animals that survive to pupation. Number of animals scored indicated as number in each bar of the histogram.

